# Human antibody cocktail deploys multiple functions to confer pan-ebolavirus protection

**DOI:** 10.1101/395525

**Authors:** Anna Z. Wec, Zachary A. Bornholdt, Shihua He, Andrew S. Herbert, Eileen Goodwin, Ariel S. Wirchnianski, Bronwyn M. Gunn, Zirui Zhang, Wenjun Zhu, Guodong Liu, Dafna M. Abelson, Crystal L. Moyer, Rohit K. Jangra, Rebekah M. James, Russell R. Bakken, Natasha Bohorova, Ognian Bohorov, Do H. Kim, Michael H. Pauly, Jesus Velasco, Robert H. Bortz, Kevin J. Whaley, Tracey Goldstein, Simon J. Anthony, Galit Alter, Laura M. Walker, John M. Dye, Larry Zeitlin, Xiangguo Qiu, Kartik Chandran

## Abstract

During the unprecedented 2013–2016 Ebola virus disease (EVD) epidemic in Western Africa and in its aftermath, the passive administration of monoclonal antibodies (mAbs) emerged as a promising treatment approach^1–7^. However, all antibody-based therapeutics currently in advanced development are specific for a single member of the *Ebolavirus* genus, Ebola virus (EBOV), and ineffective against divergent outbreak-causing ebolaviruses, including Bundibugyo virus (BDBV) and Sudan virus (SUDV)^2,3,5,7^. Here we advance MBP134, a cocktail of two broadly neutralizing human mAbs targeting the filovirus surface glycoprotein, GP, as a candidate pan-ebolavirus therapeutic. One component of this cocktail is a pan-ebolavirus neutralizing mAb, ADI-15878, isolated from a human EVD survivor^8,9^. The second, ADI-23774, was derived by affinity maturation of a human mAb^8,9^ via yeast display to enhance its potency against SUDV. MBP134 afforded exceptionally potent pan-ebolavirus neutralization *in vitro* and demonstrated greater protective efficacy than ADI-15878 alone in the guinea pig model of lethal EBOV challenge. A second-generation cocktail, MBP134^AF^, engineered to effectively harness natural killer (NK) cells afforded additional, unprecedented improvements in protective efficacy against EBOV and SUDV in guinea pigs relative to both its precursor and to any mAbs or mAb cocktails tested previously. MBP134^AF^ is a best-in-class mAb cocktail suitable for evaluation as a pan-ebolavirus therapeutic in nonhuman primates.

---

We previously isolated and characterized 349 GP-specific mAbs from a survivor of the West African EVD epidemic^9^. A systematic analysis of this library for breadth of the neutralizing mAb response against ebolaviruses identified ADI-15878 as a promising candidate therapeutic^8^. ADI-15878 possesses potent pan-ebolavirus neutralizing activity through its recognition of a highly conserved conformational fusion-loop epitope in GP with subnanomolar affinity and enhanced targeting of a cleaved GP intermediate generated in late endosomes. *In vivo*, ADI-15878 fully protected mice challenged with EBOV and SUDV and partially protected ferrets challenged with BDBV^8,9^.

Because previous work has demonstrated the predictive value of the guinea pig models of ebolavirus challenge for therapeutic antibody efficacy in nonhuman primates^10–12^, we exposed guinea pigs to a uniformly lethal dose of guinea pig-adapted EBOV (EBOV-GPA)^13^ and then treated them with equivalent doses of ADI-15878 or the ZMapp cocktail^2^ at 3 days post-challenge. Monotherapy with ADI-15878 afforded only 40–50% survival, and was less effective than ZMapp (70–100% survival), supporting prior observations that mAb cocktails targeting multiple GP epitopes may be necessary for complete protection (**Fig. 1A–B**)^1,2,8,14–16^. Accordingly, we sought to identify one or more suitable partner mAb(s) for ADI-15878. Potential cocktail components were filtered on the basis of: (i) their host origin (human or nonhuman primate); (ii) neutralization potency (50% inhibition of EBOV infectivity [IC_50_] <10 nM); (iii) anti-ebolavirus binding and neutralization breadth (recognition and neutralization of EBOV, BDBV, and SUDV); (iv) neutralization of both uncleaved and endosomally cleaved forms of ebolavirus GP, a feature previously associated with antiviral potency and breadth^8,16^; (v) lack of cross-reactivity with the secreted glycoprotein sGP present at high levels in EVD patients (to prevent sGP from “soaking up” circulating mAbs)^17,18^; and (vi) demonstrated post-exposure protection against EBOV in mice. This selection process yielded two candidates, ADI-15946 ^8,9^ and CA45^16^. Like ADI-15878, these broadly neutralizing mAbs recognize conserved conformational epitopes in the “base” subdomain of the trimeric GP spike (**Fig. 1C**)^8,16^, raising the possibility that they can compete with ADI-15878 for GP binding and limit cocktail efficiency. Indeed, preincubation of EBOV GP with saturating amounts of ADI-15878 abolished binding by CA45 in a biolayer interferometry assay (**Fig. 1D**) but did not interfere with ADI-15946 binding (**Fig. 1E**), indicating that ADI-15878 and CA45 compete for GP recognition, whereas ADI-15878 and ADI-15946 do not. Therefore, we selected ADI-15946 for further evaluation as a cocktail partner for ADI-15878.

**Figure 1.**
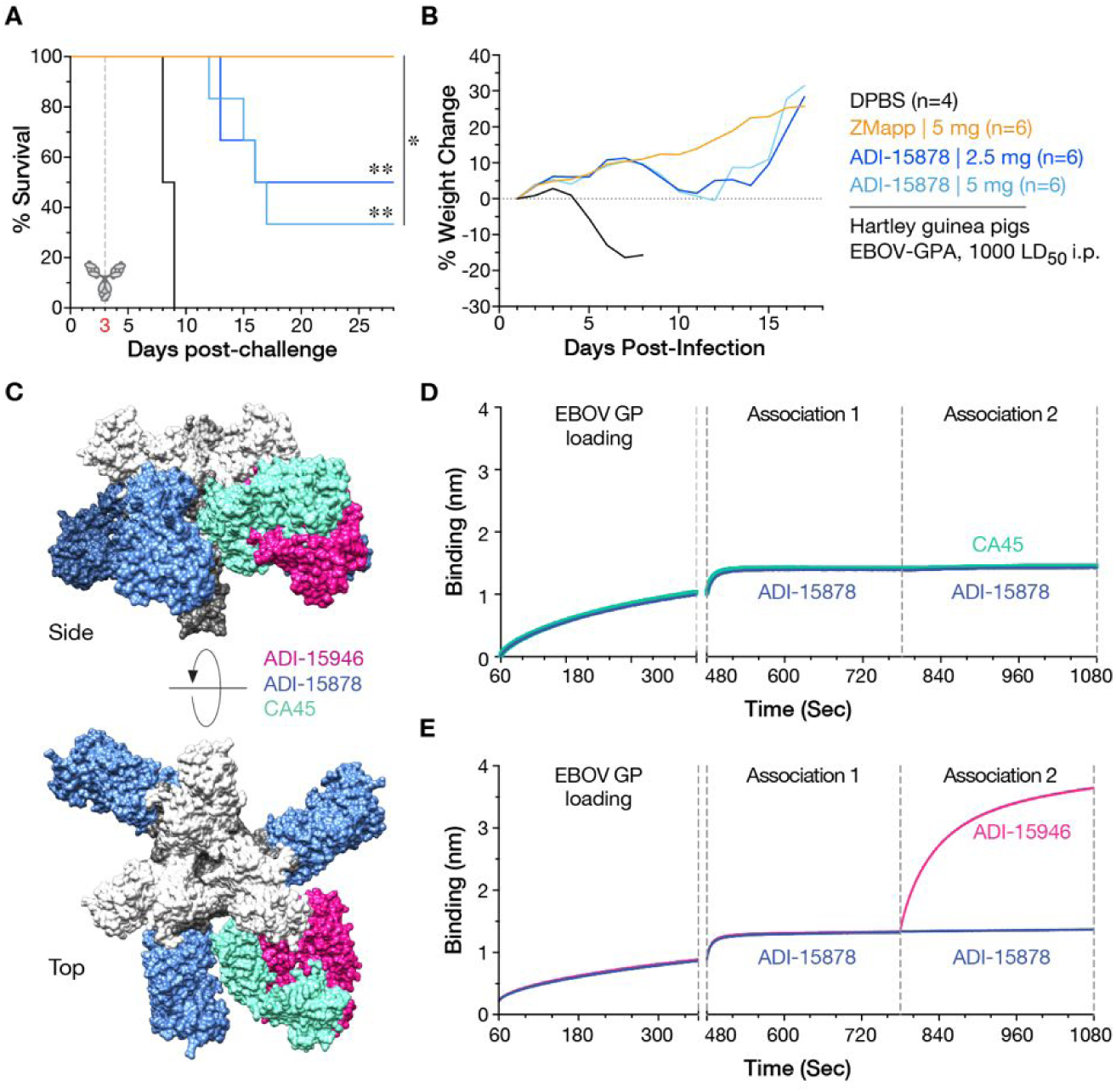
Selection of ADI-15946 as a candidate cocktail partner for ADI-15878. **A**, Hartley guinea pigs were challenged with guinea-pig adapted EBOV (EBOV-GPA) and then treated with single doses of the ZMapp cocktail or ADI-15878 or vehicle (Dulbecco’s phosphate-buffered saline, DPBS) at three days post-exposure. **, P<0.01. *, P<0.05. **B**, Combined body weights of surviving animals in each treatment group in panel A. Data from single cohorts are shown. **C**, *In silico* models of the ADI-15878, ADI-15946, and CA45 Fabs were fitted into negative-stain EM reconstructions of GP:Fab complexes^8,16^ and are superimposed onto a single EBOV GP structure [PDB ID: 5JQ3^37^] to illustrate the approximate binding footprint and angle of approach of ADI-15946 and CA45 relative to ADI-15878. **D–E**, Analysis of competitive binding of candidate IgGs to EBOV GP by biolayer interferometry. Each GP-bearing probe was sequentially dipped in analyte solutions containing ADI-15878 and then ADI-15878 (**D–E**, control), CA45 (**D**), or ADI-15946 (**E**). Results from a representative experiment are shown.

We previously showed that ADI-15946 neutralizes EBOV, BDBV, and SUDV; however, it is less potent against SUDV^8^ (see **Fig. 3A**). This neutralization deficit arises from the reduced binding affinity of ADI-15946 for SUDV GP relative to EBOV GP (**Figs. 2A and Extended Data Fig 2A**), consistent with the sequence divergence between these glycoproteins at the proposed sites of mAb contact^8^. To ensure that our mAb cocktail effectively targets all known virulent ebolaviruses at two distinct sites, we sought to specificity-mature ADI-15946 to SUDV GP using yeast-display technology^19^. Libraries were generated by introducing diversity into the ADI-15946 heavy and light chains (HC and LC, respectively) through oligonucleotide-based mutagenesis and transformation into *Saccharomyces cerevisiae* by homologous recombination. Improved variants were identified after 2 (LC) or 3 (HC) rounds of selection with a recombinant SUDV GP protein and cross-screening for retention of EBOV and BDBV binding (**Extended Data Fig. 1**). Combining beneficial LC and HC mutations yielded a variant, ADI-23774, with 5–10X enhanced binding affinity to SUDV GP and slightly improved binding to EBOV and BDBV GP relative to its ADI-15946 parent (**Fig. 2 and Extended Data Fig. 2A–C**). These gains in GP:mAb affinity effected by specificity maturation were primarily driven by reductions in the dissociation rate constant (k_off_) (**Fig. 2C and Extended Data Fig. 2D**). Next, because *in vitro* affinity maturation can increase antibody polyspecificity with potential risks of off-target binding and reduced serum half-life *in vivo*^*20*^, we assessed the polyspecificity of ADI-15946 and ADI-23774 as described^19,21^. Fortuitously, specificity maturation also reduced ADI-23774’s nonspecific binding relative to that of ADI-15946 (**Fig. 2D**). Thus, both ADI-15878 and ADI-23774 display a low level of polyspecificity, a highly desirable property for early-stage therapeutic candidates.

**Figure 2.**
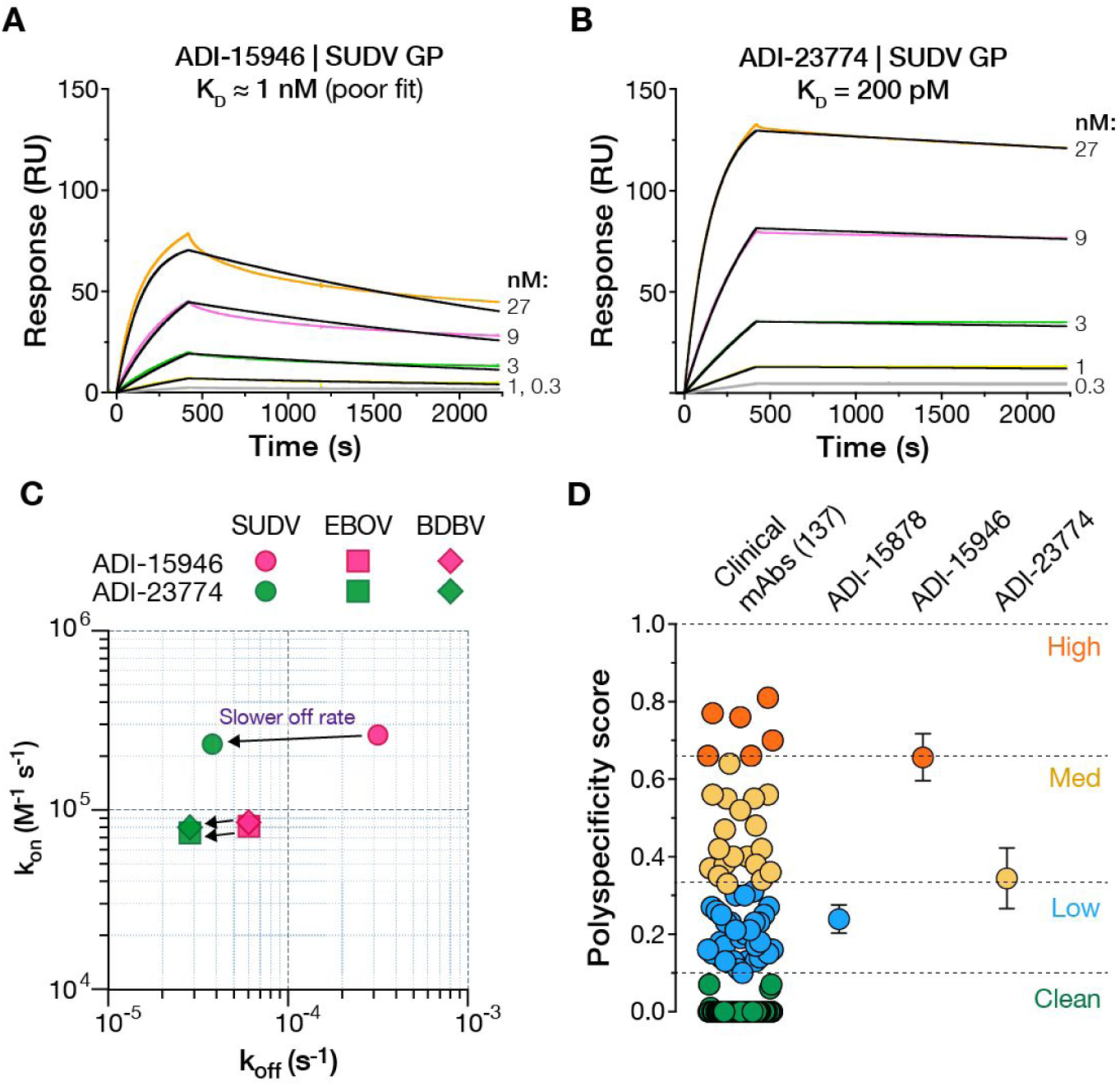
Binding and polyspecificity properties of ADI-15946 and its specificity-matured variant ADI-23774. **A–B**, Biolayer interferometry (BLI) sensorgrams for IgG-SUDV GP interactions with ADI-15946 (**A**) and ADI-23774 (**B**). Experimental curves (colored traces) were fit using a 1:1 binding model (black traces). The corresponding flow analyte (GP) concentration is indicated at the right of each curve. See **Extended Data Fig. 2D** for kinetic binding constants derived from these fits. **C**, Comparison of association (k_on_) and dissociation (k_off_) rate constants for IgG interactions with EBOV, BDBV, and SUDV GP. Arrows indicate changes in the values of these constants following ADI-15946 specificity maturation. **D**, Polyspecificity scores for candidate mAbs were determined as described previously^19^. The scores for 137 mAbs in commercial clinical development^21^ are shown for comparison. Averages±SD (n=3 for ADI-15878 and ADI-15946, n=2 for ADI-23774) from 2–3 independent experiments.

**Figure 3.**
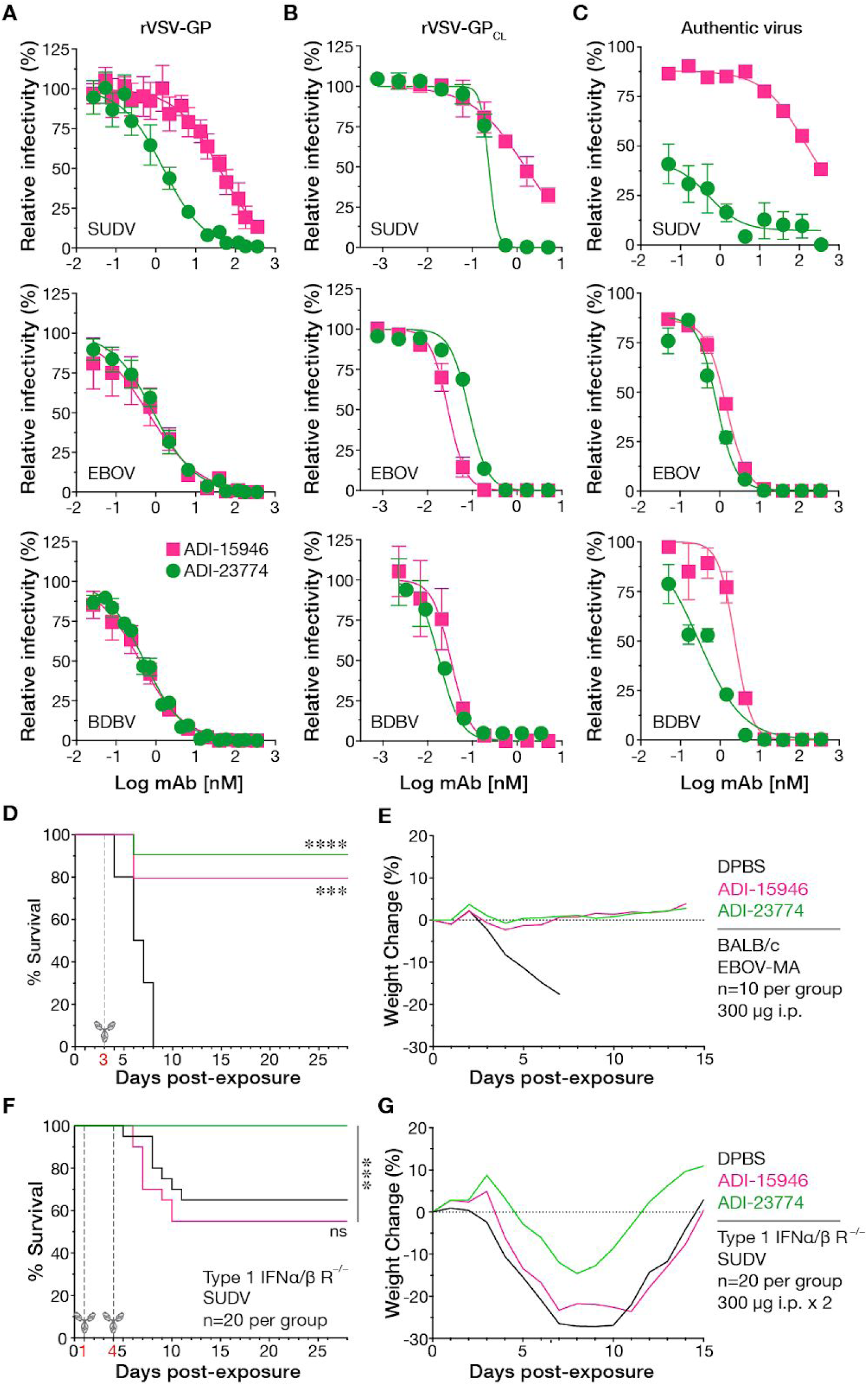
Neutralizing activity of ADI-23774 and its protective efficacy in mice. **A–B**, Neutralization of rVSVs encoding enhanced green fluorescent protein (eGFP) and bearing uncleaved (**A**) or cleaved (**B**) ebolavirus GP proteins (rVSV-GP and rVSV-GP_CL_, respectively) in Vero grivet monkey cells. Virions were preincubated with increasing concentrations of each mAb and then exposed to cells for 12 to 14 hours at 37°C. Infection was measured by automated counting of eGFP+ cells and normalized to infection obtained in the absence of antibody. Averages±SD (n=6–9 in panel A, n=3 in panel B) from 3 independent experiments. **C**, Neutralization of authentic filoviruses in Vero E6 cells measured in a microneutralization assay. Virions were preincubated with increasing concentrations of each mAb and then exposed to cells for 48 h at 37°C. Infected cells were immunostained for viral antigen and enumerated by automated fluorescence microscopy. Averages±SD (n=3) from two independent experiments. **D**, BALB/c mice were challenged with mouse-adapted EBOV (EBOV-MA) and then treated with single doses of the indicated mAbs or vehicle (DPBS) at 3 days post-exposure. **E**, Combined body weights of surviving animals in each treatment group in panel D. Data from single cohorts are shown. **F**, Type 1 IFNα/β R^−/-^ mice were challenged with wild-type SUDV and then treated with two doses of the indicated mAbs or vehicle (DPBS) at 1 and 4 days post-exposure. **G**, Combined body weights of surviving animals in each treatment group in panel F. Data pooled from two cohorts are shown. ***, P<0.001. ****, P<0.0001.

Concordant with ADI-23774’s improved binding to both recombinant GP proteins (**Fig. 2**) and rVSVs bearing full-length transmembrane GP (**Extended Data Fig. 2A**), it neutralized SUDV more potently than ADI-15946 [mAb concentration at half-maximal neutralization, IC_50_ (±95% confidence intervals) = 1±1 nM for ADI-23774 vs. 35±1 nM for ADI-15946]. Further, unlike ADI-15946, ADI-23774 could neutralize rVSV-SUDV GP completely without leaving an un-neutralized fraction (**Fig. 3A, C**). These trends were also apparent against viruses bearing a proteolytically cleaved form of GP resembling an endosomal entry intermediate (**Fig. 3B**). ADI-23774’s increased effectiveness against SUDV did not compromise its capacity to neutralize EBOV and BDBV (**Fig. 3A, C**).

To assess the effects of these gains in neutralization potency on mAb protective efficacy, we evaluated ADI-23774 in murine models of EBOV^22^ and SUDV^23^ challenge. ADI-23774 resembled ADI-15946 in its capacity to protect animals from challenge with mouse-adapted EBOV when administered 3 days post-exposure (**Fig. 3D–E**). However, unlike ADI-15946, which conferred no benefit against SUDV^8^, ADI-23774 stemmed weight loss in SUDV-challenged mice and fully protected them when administered 1 and 4 days post-exposure (**Fig. 3F–G**). ADI-23774 is an optimized candidate to serve as a partner mAb for ADI-15878, with enhancements over its precursor in biophysical properties, neutralization breadth, and anti-ebolavirus protective breadth in mice.

We combined ADI-15878 and ADI-23774 into a cocktail, MBP134, and examined its functional properties *in vitro*. MBP134 potently neutralized recombinant vesicular stomatitis viruses (rVSVs) bearing EBOV, BDBV, and SUDV GP (half-maximal inhibitory concentration, IC_50_<1 nM) (**Extended Data Fig. 3A–C**). Moreover, MBP134 also neutralized an rVSV-GP bearing the spike glycoprotein from a novel, divergent ebolavirus, Bombali virus (BOMV), recently discovered in free-tailed bats in Sierra Leone^24^ (**Extended Data Fig. 3D**). These findings indicate that MBP134’s broad anti-ebolavirus coverage could extend beyond the currently recognized disease-causing agents to unknown viruses and viral variants that may pose a spillover risk in the future.

To evaluate the protective potential of MBP134, we exposed guinea pigs to a uniformly lethal dose of EBOV-GPA and then treated them with increasing doses of MBP134 at 3 days post-challenge (**Fig. 4A–B**). Treatment with 2.5–3.3 mg of MBP134 (total mAb dose) per animal afforded near complete protection (**Fig. 4A**), whereas treatment with ADI-15878 alone at 2.5–5 mg per animal afforded ≤50% protection (**Fig. 1A**), providing evidence for the superior efficacy of the cocktail over the monotherapy.

**Figure 4.**
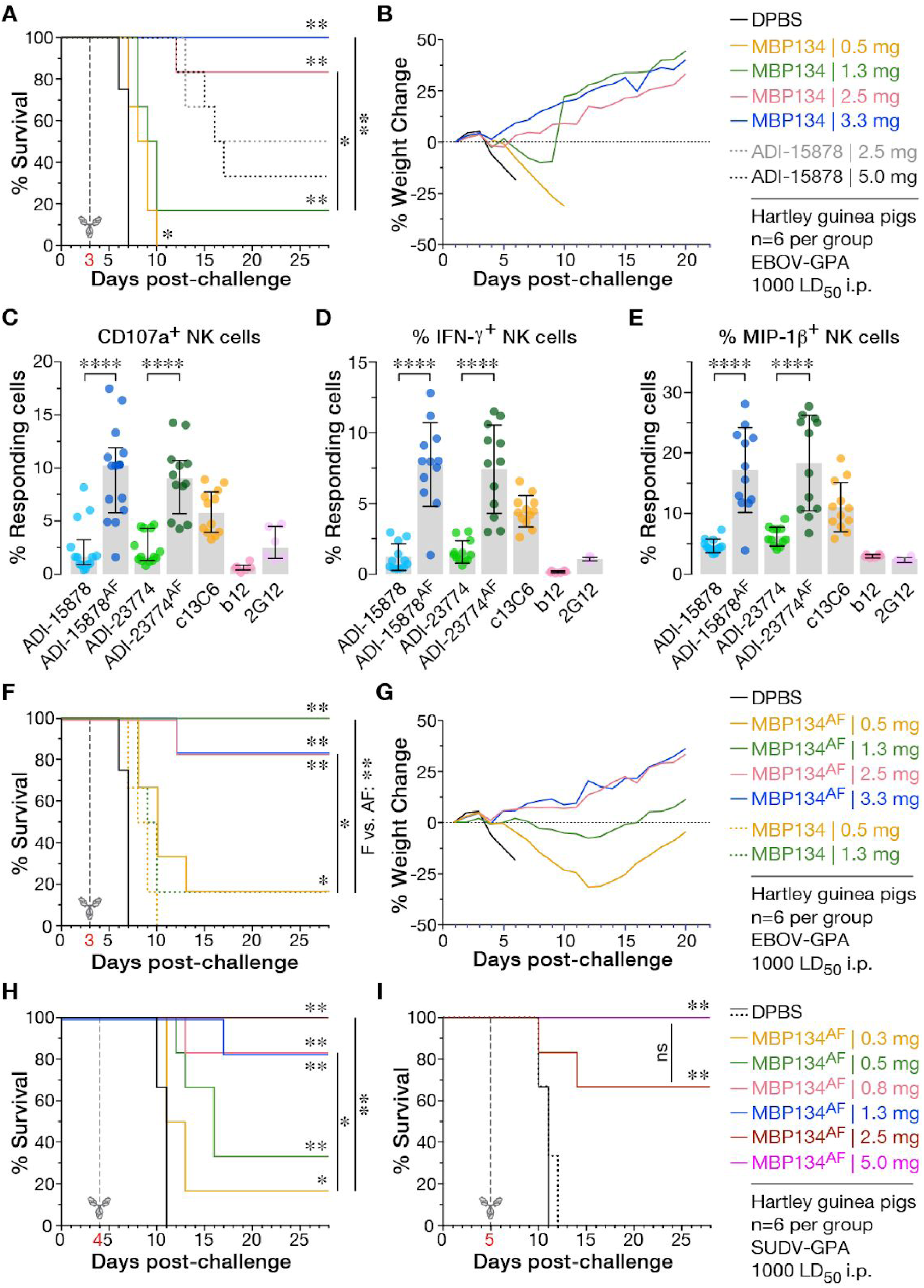
Protective efficacy of MBP134 and development and evaluation of the second-generation MBP134^AF^ cocktail. **A–B**, Hartley guinea pigs were challenged with EBOV-GPA and then treated with single doses of ADI-15878, MBP134 (1:1 mixture of ADI-15878 and ADI-23774), or vehicle (DPBS) at three days post-exposure. **B**, Combined body weights of surviving animals in each treatment group in panel A. Data from single cohorts are shown. **C–E**, Activation of human natural killer (NK) cells by EBOV GP:IgG complexes. NK cells enriched from the peripheral blood of four different human donors were incubated with complexes between EBOV GP and the indicated IgGs (10 µg/mL) for 5 h at 37°C, and then stained with antibody-fluorophore conjugates specific for the cell-surface markers CD3, CD56, and CD16, followed by intracellular staining for markers of NK cell activation, CD107a (degranulation) (**C**), IFN-γ (**D**), and MIP-1β (**E**). CD3^−^/CD56^dim^/CD16^+^ NK cells were analyzed by flow cytometry. Data with cells from all four donors are pooled. HIV-1 glycoprotein-specific mAbs b12 and 2G12 are included as (negative) controls for antigen specificity. EBOV GP-specific mAb c13C6 produced in transgenic *N. benthamiana* tobacco plants to bear a highly functional afucosylated/agalactosylated bisected glycan is included as a positive control. Averages±SD (n=12–14 for all mAbs except b12 and 2G12 (n=6) from 4 independent experiments. **F**, Hartley guinea pigs were challenged with EBOV-GPA and then treated with single doses of MBP134, MBP134^AF^ (1:1 mixture of ADI-15878^AF^ and ADI-23774^AF^), or vehicle (DPBS) at 3 days post-exposure. **G**, Combined body weights of surviving animals in each treatment group in panel F. Data are from single cohorts. **H–I**, Guinea pigs were challenged with SUDV-GPA and then treated with single doses of MBP134^AF^ or vehicle (DPBS) at 4 (**H**) or 5 (**I**) days post-exposure. Data from single cohorts are shown. **, P<0.01. *, P<0.05. ns, P>0.05.

To further enhance the *in vivo* potency of MBP134, we set out to optimize an orthogonal property of its component mAbs—their capacity to harness the host antiviral innate immune response through antigen-dependent engagement of activating Fc receptors on immune cells. Because cytolysis by natural killer (NK) cells has been demonstrated to play a critical role in protection against EBOV infection^25^, we measured the EBOV GP-dependent activation of human NK cells derived from four seronegative human donors by ADI-15878 and ADI-23774. Both mAbs were poorly active, as judged by their limited capacity to induce three markers of NK cell activation—degranulation (**Fig. 4C**), and production of IFN-γ (**Fig. 4D**) and MIP-1β (**Fig. 4E**)—relative to the ZMapp component mAb c13C6^2^. Thus, viral neutralization appears to be the primary mode of MBP134’s antiviral activity. We postulated that NK cell recruitment, activation, and infected-cell killing by MBP134 could be improved by enhancing its capacity to engage the activating Fc receptor FcγRIIIa on NK cells. Indeed, previous work has demonstrated that mAbs bearing uniformly afucosylated glycan structures display precisely this property^26,27^, which has been linked to their enhanced protective efficacy against several viral pathogens, including EBOV^28–30^. Accordingly, we generated and evaluated afucosylated variants of the MBP134 mAbs (ADI-15878^AF^ and ADI-23774^AF^, respectively). Both variant mAbs significantly outperformed their fucosylated precursors in all three NK cell activation assays (**Fig. 4C–E**), confirming their enhanced capacity to engage the innate immune system.

Finally, we assessed the protective potential of the second-generation MBP134^AF^ cocktail in guinea pigs, which possess an Fc receptor (gpFcγRIV) homologous to human FcγRIIIA and with enhanced binding affinity for afucosylated human IgGs relative to their fucosylated counterparts^31^. Only 1.3 mg (total mAb dose) of MBP134^AF^ administered at 3 days post-challenge was required to fully protect animals from EBOV-GPA (**Fig. 4F–G**)—an approximately twofold reduction in dosage relative to the parent MBP134 cocktail (P=0.0064) (**Fig. 4A**). Moreover, a single MBP134^AF^ dose as low as 0.8 mg administered at 4 days post-challenge was sufficient to fully protect guinea pigs challenged with guinea pig-adapted SUDV (SUDV-GPA)^32^ (**Fig. 4H**). Remarkably, MBP134^AF^ afforded 70–100% protection from SUDV-GPA challenge (at 2.5–5.0 mg total mAb dose) even when treatment was delayed to 5 days post-challenge (**Fig. 4I**). These are, to our knowledge, the lowest single doses (0.8–1.3 mg total mAb) and longest post-exposure treatment windows (4–5 days) demonstrated to protect guinea pigs from lethal ebolavirus challenge. All previous studies with both therapeutic mAbs and mAb cocktails used total doses of 5–10 mg and did not report initial mAb treatments past 3 days post-exposure^15,16,33–36^.

Herein, we have used a rigorous mAb selection strategy coupled to yeast-based specificity maturation and Fc glycan engineering to develop a next-generation human mAb cocktail, MBP134^AF^, that can broadly protect against disease-causing ebolaviruses in small and large rodents. MBP134^AF^ targets all known ebolavirus glycoproteins (including that from a recently discovered “pre-emergent” agent) at two independent antigenic sites and neutralizes both their extracellular and endosomal intermediate forms, thus conferring potency as well as robustness to viral neutralization escape. Further, it is optimized to leverage both mechanical neutralization and Fc-linked innate immune functions to block viral infection and spread. By virtue of these advanced features, MBP134^AF^ affords unparalleled improvements in therapeutic potency against broad ebolavirus challenge in the stringent guinea pig challenge model relative to any mAbs or mAb cocktails that have been described previously. Our findings set the stage for evaluation of this best-in-class mAb cocktail as a pan-ebolavirus therapeutic in nonhuman primates (see companion manuscript, Bornholdt *et al.*).

## Acknowledgments

We thank Tyler Krause, Cecelia Harold, Tanwee Alkutkar, and Isabel Gutierrez for technical support. We thank M. Javad Aman for provision of mAb CA45. This work is supported by NIH grants AI1332204 and AI132256 (to K.C.) and U19AI109762 (to KC., J.M.D., and L.Z.), and by the Public Health Agency of Canada (to X.Q.). K.C. was additionally supported by an Irma T. Hirschl/Monique Weill-Caulier Research Award. Opinions, conclusions, interpretations, and recommendations are those of the authors and are not necessarily endorsed by the US. Army. The mention of trade names or commercial products does not constitute endorsement or recommendation for use by the Department of the Army or the Department of Defense. L.Z., X.Q., D.M.A., and Z.A.B. were supported by DTRA contract HDTRA1-13-C-0018.

## Author Contributions

K.C., Z.A.B., L.Z., L.M.W., X.Q., and J.M.D. conceived the study. E.G. and L.M.W. performed the specificity maturation of ADI-15946. D.M.A and Z.A.B. expressed and purified mAbs and GP ectodomain proteins. A.Z.W. and L.M.W. carried out binding studies by surface plasmon resonance, and Z.A.B. performed competition binding studies by biolayer interferometry A.Z.W. performed binding studies by ELISA. A.Z.W., A.S.W., R.K.J., and R.B. carried out VSV-based neutralization experiments. A.S.H., R.M.J., and R.R.B. performed authentic virus neutralization experiments. B.G. and G.A. carried out mAb effector function studies. A.S.H. performed the mouse challenge studies. S.H., Z.Z., W.Z., G. L. and X.Q. performed the guinea pig challenge studies. T.G. and S.J.A. provided key BOMV reagents. K.C., A.Z.W. and Z.A.B. wrote the manuscript with contributions from all of the authors.

## Supplementary Materials

Methods

Extended Data Figures 1–3

